# Frame-by-frame annotation of video recordings using deep neural networks

**DOI:** 10.1101/2020.06.29.177261

**Authors:** Alexander M. Conway, Ian N. Durbach, Alistair McInnes, Robert N. Harris

## Abstract

Video data are widely collected in ecological studies but manual annotation is a challenging and time-consuming task, and has become a bottleneck for scientific research. Classification models based on convolutional neural networks (CNNs) have proved successful in annotating images, but few applications have extended these to video classification. We demonstrate an approach that combines a standard CNN summarizing each video frame with a recurrent neural network (RNN) that models the temporal component of video. The approach is illustrated using two datasets: one collected by static video cameras detecting seal activity inside coastal salmon nets, and another collected by animal-borne cameras deployed on African penguins, used to classify behaviour. The combined RNN-CNN led to a relative improvement in test set classification accuracy over an image-only model of 25% for penguins (80% to 85%), and substantially improved classification precision or recall for four of six behaviour classes (12–17%). Image-only and video models classified seal activity with equally high accuracy (90%). Temporal patterns related to movement provide valuable information about animal behaviour, and classifiers benefit from including these explicitly. We recommend the inclusion of temporal information whenever manual inspection suggests that movement is predictive of class membership.

## 1 Introduction

Technological advances in quality, size, battery life and storage capacity have enabled video cameras to record more data at better quality on a broader variety of animals, becoming small enough to deploy on numerous animal species (Rutz & Troscianko, 2013; Takahashi et al., 2004) and on drones (Anderson & Gaston, 2013; Cruzan et al., 2016), as well as in more conventional fixed locations. Footage captured using video cameras needs to be annotated for use in scientific research, a currently labour intensive process often involving highly trained scientists manually annotating the content of videos frame by frame. Even with dedicated annotation software, this presents a major bottleneck for scientific research based on these data, necessitating the development of computer-assisted approaches (Schneider, Taylor, Linquist, & Kremer, 2019; Weinstein, 2015).

Video classification is a challenging modelling problem, with the challenges of image classification amplified because the same sources of natural visual variation occur not only between videos but also within videos as objects move around and change poses, scales, illuminations and backgrounds during the course of a single video. The video camera itself can move around during recording, introducing additional variation, particularly in environments where cameras move due to wind or water movement, or because cameras are attached to animals moving around their environment. The temporal component of video also presents significant modeling challenges not only because it dramatically increases the size of video data but because the relevant visual features required to classify a video can span several frames with no single frame containing enough information on its own. The pixels of an image representing objects are not only correlated spatially to form visual object features in a single frame but are also correlated through time. Like image classification, traditional computer-based approaches to video classification have primarily used feature engineering algorithms that create input variables based on predetermined traits. Spatial algorithms construct variables such as Harris or SIFT features (Lowe, 2004) that discriminate patterns within an image (e.g. morphometric features), while spatio-temporal algorithms such as the Cuboid and Harris-3D detectors (Dollár, Rabaud, Cottrell, & Belongie, 2005) capture additional motion information between frames. The main limitations of these approaches arise from their need to know how to represent input features in advance – this requires substantial knowledge of the study species, and hinders generalization across species and environmental contexts (Schneider et al., 2019).

Deep neural networks (DNNs) are highly flexible machine learning models that use stacked nonlinear combinations of inputs together with a gradient descent learning procedure to jointly learn feature representations together with how these should be translated into classifications, based on labeled data, thus avoiding the main drawback of feature engineering. DNNs are the current state-of-the-art for many challenging perceptual problems involving image, video, audio or text, where hand-designing input feature representations is nontrivial (Liu, Wang, Liu, Liu, & Alsaadi, 2016).

Convolutional neural networks (CNNs) are a specialized kind of DNN architecture that takes advantage of the characteristics of image data to learn hierarchies of local features that are invariant to common translation operations like shifting, stretching and rotation. This reduces the number of required parameters while leaving enough representational power to achieve good performance on image classification and other tasks involving data that have a regular grid-like topology of locally correlated hierarchical features. CNNs typically involve a stacked sequence of convolutional layers – traversing the network, the output of each of these layers can be thought of as an increasingly complex summary or ‘encoding’ of the input image as a one-dimensional numeric vector. CNNs have found numerous, and increasing, applications in ecological studies (Christin, Hervet, & Lecomte, 2019; Weinstein, 2018a), where image classification has been used for species identification (Gomez Villa, Salazar, & Vargas, 2017; Weinstein, 2018b; Zhang, He, Cao, & Cao, 2016), count surveys (Borowicz et al., 2018; Gray, Fleishman, et al., 2019; Torney et al., 2019), individual animal re-identification (Schneider et al., 2019), and morphometric measurement (Gray, Bierlich, et al., 2019). Applications to video classification, however, remain rare. With the exception of Trinh, Yoshihashi, Kawakami, Iida, and Naemura (2016), who combined neural network architectures to detect birds flying into wind turbines from sequences of input frames, most studies have either classified frames in isolation (Siddiqui et al., 2018), or used previous frames primarily to improve the discrimination of the focal animal from background scenery, using motion-detection algorithms (Weinstein, 2018b; Zhang et al., 2016).

There are three approaches to using DNNs for video classification beyond treating the problem as an image classification task by modeling frames independently. The simplest approach concatenates the vector encodings obtained from each of a sequence of input images to predict the class of the last image in the sequence; images in the input sequence are considered to be independent. The second approach uses the sequence of vector encodings produced from the sequence of input images as input to a second model – a recurrent neural network (RNN), a specialized architecture often used to process sequential data involving a temporal component (Donahue et al., 2014; Trinh et al., 2016). Finally, CNNs can be directly modified to incorporate motion information in videos by extending their convolution from two spatial dimensions (width and height) to three spatio-temporal dimensions (width, height and time), parameters of which are jointly estimated (Tran, Bourdev, Fergus, Torresani, & Paluri, 2015).

In this paper we have used these approaches to perform frame-by-frame annotation of two video datasets. The first was taken from a fixed underwater camera placed inside nets at a salmon trap net fishery in Scotland, for the purpose of detecting seal visits to salmon nets and ultimately reducing conflict between fisheries and seals. Here the task was to detect whether a seal is present in a frame, based on that and preceding frames. The second dataset was collected by animal-borne cameras deployed on African penguins in South Africa. Here the purpose was to replicate manual annotations allocating each frame to one of six pre-defined classes covering diving and surface behaviour exhibited by the birds. To the best of our knowledge, this is the first time DNNs have been applied to annotate animal-borne video. For each dataset, our primary goal was to evaluate whether incorporating the temporal component of video brings any improvement in classification accuracy, relative to an image-only benchmark.

## 2 Materials and Methods

### 2.1 Data

#### 2.1.1 Seals

An underwater video system was used to study seal behaviour at a salmon trap net fishery in north east Scotland in 2015 as part of a programme of research aimed at reducing conflict between fisheries and seals. Cameras were placed inside static coastal nets to monitor seals as they moved in and out of nets to depredate salmon. There was no artificial lighting and so the cameras recorded during hours of daylight.

The labelled component of the dataset consisted of six video recordings of ca 140 minutes each, converted into images at 4fps. A total of 152 instances in which a seal entered the net were observed by manual inspection, and entry and exit times for each of these recorded (Figure A.1, Appendix A). Visits lasted between 2s and 59s, with an average duration of 13.5s. Seals were not visible in frame for the entire duration of a visit, so all images between the start and end times of a recorded visit were manually inspected and labelled as containing a seal or not. After processing, there were 4419 images containing a seal. While the vast majority of footage does not contain a seal in frame, we restricted the number of absence images to 7809, roughly twice the number of seal images, to avoid a large class imbalance. Absence images were collected by randomly sampling segments of video from the remainder of the video. Images from four videos were used to train models (3826 seal, 6949 no seal), while images from each of the remaining two videos were used as validation (407 seal, 973 no seal) and test (192 seal, 111 no seal) datasets respectively.

#### 2.1.2 Penguins

Animal-borne video recorders (AVR) were deployed on breeding African penguins attending small chicks at Stony Point, South Africa, between 2015 and 2016 (McInnes, McGeorge, Ginsberg, Pichegru, & Pistorius, 2017). The AVRs were tube-shaped, and together with the casing weighed 100g with dimensions 104×26×28mm. Devices were attached to the lower backs of the penguins with strips of waterproof tape during the evening preceding an anticipated foraging trip. AVRs were programmed to divide the battery life into two recording bins of ca 30 min each, at sunset and midday to reflect potential temporal differences in diving behaviour. Recorders where retrieved when the bird returned to the colony, either on the same day that the bird was at sea and after the bird had time to provision its chicks, between 16:00 and 20:00, or the following morning if the bird could not be located the previous day.

The labelled component of the dataset consisted of 12 video recordings of ca 30 minutes each, again converted into images at 4fps. These were manually classified into five diving behaviours (subsurface diving (less than 1m); shallow diving (1-5 m); and the descent, bottom, and ascent phases of deep dives) and one surface behaviour (searching, see Figure A.2, Appendix A). A total of 52722 images were obtained, with substantial imbalance between behaviours (Table A.1, Appendix A). Images from nine videos were used to train models (41958 images, see Table A.1 for distribution over behaviours), while images from the remaining videos were used as validation (two videos, 7168 images) and test (one video, 3596 images) datasets respectively.

### 2.2 Neural networks

We consider four broad classes of models, of increasing complexity. The first ignores the temporal aspect of video data and attempts to classify each image independently using a standard CNN-based approach. Pretrained CNNs (VGG16, ResNet50, Inception v3 and Inception-ResNet v2) were truncated at an intermediate layer – the output of this intermediate layer summarizes or ‘encodes’ an image in a one-dimensional vector. Up to three dense layers were added to the truncated network, and a new output layer added for the (seal or penguin) classification task. The second model used the same approach, but classified an image by first concatenating the vector encoding obtained from the truncated layer for that image with similar vectors obtained for the previous *F* − 1 images. This concatenated vector, which summarizes a set of *F* consecutive images rather than (as in the first model) just a single image, was then passed these to subsequent dense layers as before. The third model was the spatial-then-temporal model described in the introduction. To classify a single image, it took the vector encodings from the last *F* images (including the current image), as in the previous model, but instead of concatenating the encodings it passed these as input to a recurrent neural network, which combined these temporally (Figure 1). We used two pre-trained CNNs to encode frames (ResnNet50, VGG16) and three different RNN architectures (Long Short-Term Memory (LSTM), SimpleRNN, Gated Recurrent Units (GRU)). One key step was to pre-compute the frame vector encodings from the pre-trained CNN models so that these did not have to be re-computed in each RNN model. A single training epoch for the mixed recurrent convolutional network (RCNN) architecture with a VGG encoder took approximately 15 minutes without pre-computation but only 3 seconds with pre-computed features (because most of the computation time was spent in the CNN part of RCNN). The final model jointly modelled spatial and temporal aspects using a 3-dimensional CNN that convolves simultaneously over space and time. Because convolutions occur simultaneously over space and time, the 3-D CNN cannot leverage pre-computation, and generators had to be used to stream the data from disk to avoid out-of-memory problems. Despite various attempts at optimization, a single model took approximately 3 days to converge on a single GPU, and returned substantially worse accuracy than even an image-only model. We therefore do not report on these results further. We chose model hyperparameters using a grid search over the number of nodes in each of the three dense layers in Model 1 and 2 (32, 64, 96,…, 512), the dropout rate (0, 0.1, 0.2,…, 0.5), and the length of the sequence of images used in Models 2 and 3 (1, 3, 5, 7, 9, …, 31). Following Krizhevsky, Sutskever, and Hinton (2012), each model’s weights were initialized using the Xavier initialization and each model was trained in 3 rounds of 20 epochs with an early stopping patience of 5 epochs using the Adam optimizer (Kingma & Ba, 2014). The learning rate was initially set to 0.001 and reduced by a factor of 10 between training rounds, and max pooling was used. Models were evaluated based on test set accuracy (proportion of all predictions that were correct), precision (proportion of positive predictions that were correct), and recall (proportion of positive examples correctly predicted). For the seals dataset, seal presence is a natural choice for the positive class. For multi-class classification problems, precision and recall were obtained for each class, and overall precision and recall calculated as an average of these, weighted by sample size. Models were implemented using the TensorFlow (Abadi et al., 2016) library with Keras (Chollet et al., 2015). Training and testing were done on a three separate Linux virtual machine instances running on Google Cloud Platform, each with eight Nvidia Tesla K80 Graphics Processing Units (GPUs), 160 GB of RAM and 32 CPU cores. Code and analysis scripts are available online at https://github.com/alxcnwy/Deep-Neural-Networks-for-Video-Classification.

**Figure 1.**
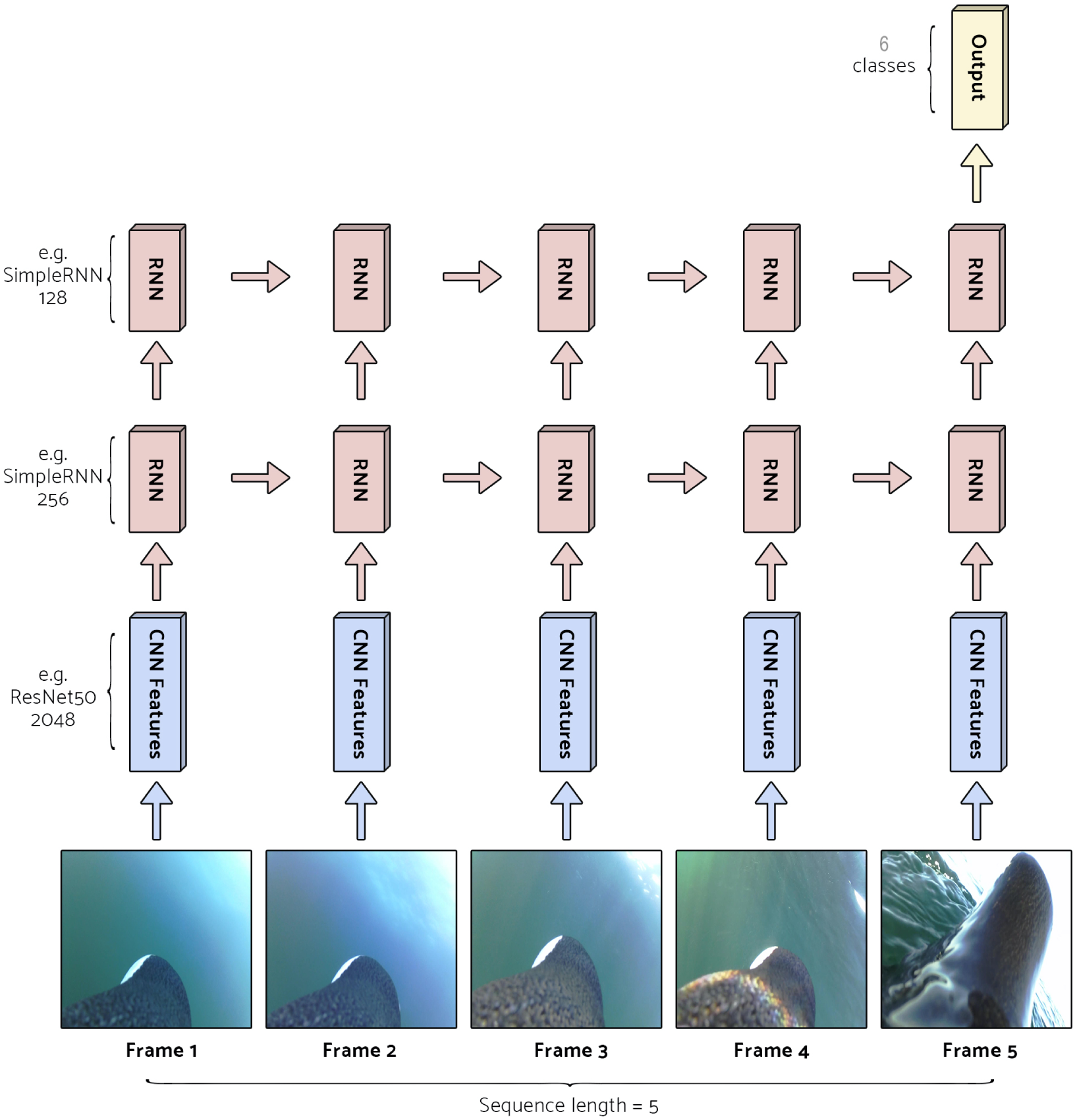
A “spatial-then-temporal” neural network for frame-by-frame video classification. To predict the class of a frame (Frame 5), a pre-trained, truncated CNN (e.g. ResNet50) is used to summarize or ‘encode’ each of a sequence of images (here, the last five frames) as one-dimensional numeric vectors. The sequence of vector encodings is then used as input in a recurrent neural network (RNN), here shown using two SimpleRNN layers. The RNN outputs predicted probabilities that the behaviour in the final frame is of type *i, i* = 1, …, 6.

## 3 Results

A video component did not bring meaningful benefits in detecting seals, with both image-only and video models accurately classifying 89% of images in the test set, and small improvements in precision being offset by marginally worse recall (Table 1). Most incorrect classifications occurred at the beginning and end of visits, as the seal was entering or exiting the field of view and where only a small part of the seal may be in view (Figure B.1, Appendix B). All 152 seal visits across training, validation, and test sets were detected by either model.

**Table 1:** Classification accuracy for three best video models and best image model. Including temporal information in the form of an RCNN led to very marginal improvement in the easier seal detection task, but gave a 25% relative improvement in the ability to discriminate penguin behaviours, largely due to improved performance at the start and end of behaviours (Figure 2). Further details on the architectures and run times of these models are given in Table B.2 and B.3, Appendix B.

| Seal detection model |  |  |  |  |
| --- | --- | --- | --- | --- |
| Architecture | RCNN | RCNN | RCNN | IMAGE |
| Accuracy (Test) | 89.4% | 89.2% | 89.1% | 89.1% |
| Precision (Test) | 100% | 99.4% | 100% | 97.6% |
| Recall (Test) | 83.9% | 83.9% | 83.3% | 84.9% |
| Accuracy (Validation) | 96.3% | 95.9% | 95.7% | 93.7% |
| Accuracy (Train) | 95.4% | 95.4% | 95.3% | 95.2% |
| Penguin behaviour classifier |  |  |  |  |
| Architecture | RCNN | RCNN | RCNN | IMAGE |
| Accuracy (Test) | 85.4% | 84.0% | 84.2% | 80.5% |
| Precision (Test) | 85.4% | 84.0% | 84.2% | 80.5% |
| Recall (Test) | 87.6% | 87.6% | 85.5% | 82.8% |
| Accuracy (Validation) | 82.6% | 82.4% | 81.0% | 81.5% |
| Accuracy (Train) | 90.0% | 88.9% | 94.4% | 88.7% |

Including temporal information in video data, in the form of spatial-then-temporal models, improved the accuracy of penguin behaviour classifications from 80.5% (image-only benchmark) to 85.4%, a 25% relative reduction in classification error (Table 1), and improved both precision and recall. Models concatenating frame encodings occupied an intermediate position between full video and image-only models. Classification accuracy improved for most penguin behaviour types (Table B.1, Appendix B), but particularly for descent and bottom dive phases (precision increasing by 17% and 14%), and for shallow and subsurface dives (recall increasing by 12% and 13%). Image-only models tended to misclassify bottom dives as descent dives, and mistook parts of the ascending and descending dive phases for shallow dives. To some extent this reflects fuzzy boundaries between behavioural classes, but temporal information resolved some of these misclassifications (Figure 2). Search activity, the sole surface behaviour and also the most prevalent class, was almost perfectly discriminated.

**Figure 2.**
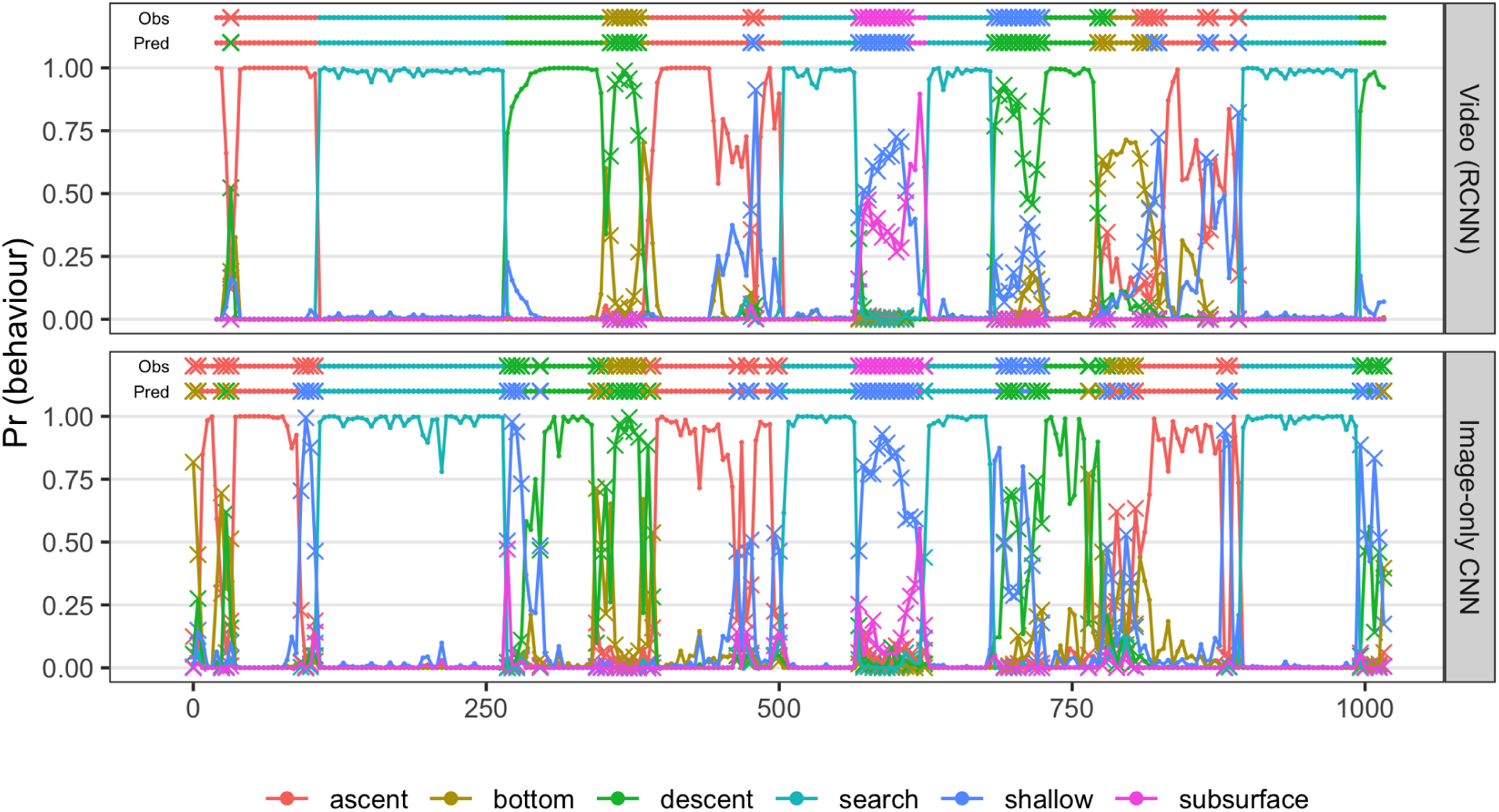
Predicted probabilities for penguin behaviour classes, with misclassifications plotted as crosses. Observed and predicted classes are plotted above the probabilities, using the same notation. Image-only models tend to misclassify bottom dives as descent dives (frame 350–390), and ascending and descending dive phases as shallow dives (frame 90–110 and 260–280). Video models resolve some of these errors. They also smooth transitions between behaviours (frame 260–280), better identify periods where classification uncertainty is high (frame 570-620, 750-850) and where alternate interpretations are possible (frame 570-620).

Preferred RCNN models for seal detection achieved a degree of parsimony by using a relatively short sequence of frames, and in exchange used relatively complex pre-trained CNN (ResNet50) and RNN (LSTM) architectures (Table B.2, Appendix B). In contrast, equivalent preferred models for penguin behaviour classification used longer sequences of frames, but simpler CNN (VGG16) and, sometimes, RNN (SimpleRNN) architectures (Table B.3, Appendix B). Both applications selected a relatively large number of nodes in the final hidden layers.

## 4 Discussion

Although images are more commonly used in ecological research and are easier to work with (Swinnen, Reijniers, Breno, & Leirs, 2014), movement information contained in video provides richer insight into animal behaviour and taking this into account can improve the identification of animals and their behaviours (Trinh et al., 2016). We found that for a relatively simple task – detecting seal activity in an image – an image-only CNN was adequate, and incorporating temporal information did not meaningfully improve out-of-sample performance, even for those difficult cases in which a seal enters or exits the field of view. For a more difficult task of inferring penguin behaviour from animal-borne cameras, using a video model led to substantial reduction in classification error over an image-only model, and was particularly useful in disentangling certain kinds of diving behaviour. In both applications accuracy is not sufficient for full automation of the tasks, but can facilitate manual processes by partially labelling the data – identifying those classes that can be accurately discriminated and pointing the researcher to segments requiring closer inspection. Our datasets were relatively small, consisting of 6-12 hours of labelled footage, and the ability of the models to generalize to new environments is unclear, but even in those classes where absolute performance was moderate, video models outperformed image-only models. Improvements are likely to be larger with larger datasets.

Practically, researchers wanting to construct a model for the frame-by-frame annotation of video have to follow a number of steps: manually labelling a subset of the data; converting the video into images; allocating these images between training, validation, and test sets; choosing appropriate neural network architectures and estimating the parameters of those models; selecting a preferred model and using it to process the unlabelled portion of the data; and linking frame-by-frame predictions to the broader research objectives for which the classifier was developed. Video data are manually annotated by recording the start and end times of events whose boundaries may be difficult to distinguish precisely. Poorly separated classes can reduce classification accuracy, and preprocessing steps for image classification sometimes remove ambiguous images to improve class separability. Video models, however, use a sequence of frames *t, t* − 1, …, *t – F* to predict the class of frame *t*, and removing ambiguous images makes the time difference between adjacent images variable. While it is possible that removing ambiguous examples may improve accuracy more than maintaining constant time difference between images, this is likely to be case-specific, and not generally recommended. Rather, the presence of ambiguous images places an effective upper limit on the accuracy that can be achieved, which may or may not impact on broader research objectives. For seal visits, for example, the detection of a seal presence is more important than identifying the exact time of entry. The first and last few frames of a visit often contain only a tiny sliver of seal or, because the times are approximate, no seal at all. These frames reduce classification accuracy but have very little bearing on the practical usefulness of the classifier.

Video data are converted to images at a user-specified frame rate, with the recording equipment setting an upper bound. A higher frame rate increases the number of images available to train models, which is always beneficial as long as there are meaningful differences between adjacent images. It is important to randomly allocate contiguous sequences of frames i.e. video sequences, to training, validation and test datasets, rather than randomly allocating the frames themselves. Doing the latter breaks apart sequences, losing potentially valuable information, and also means that very similar images occur in both training and test sets. We also recommend assessing whether the video in the test dataset has the same environmental conditions as video used to train the model (e.g. if a random segment of each file is used to test). If so, the ability of the model to generalize to new environments may be overestimated.

When building an RCNN, key choices are what frame rate and sequence length to use. These factors are study-specific, and the chosen frame rate need not be the same as the frame rate used to convert video to frames. Higher frame rates allow for fine-scale changes in movement to be captured, but the same number of frames covers a shorter time interval. Increasing sequence length requires more parameters, increasing the chances of overfitting and requiring more data. Which of the two – looking back further in time or capturing fine-scale movement – benefits classification accuracy more will be study-specific. These factors can be investigated by searching over possible frame rate/length pairs, but this quickly becomes computationally expensive. Our applications have relatively little labelled data and so we fixed the frame rate to one that would allow broad differences in behaviour, observed over a few seconds, with 5 *< F <* 10. Pretrained CNNs offer a parsimonious way of summarizing images in a form that can be passed on the second-stage RNN (Donahue et al., 2014). Our best seal model combined a relatively complex CNN and RNN with a short frame sequence, whereas the best penguin model had a simple CNN and RNN, but used a longer sequence of frames. Since model complexity is primarily achieved through more parameters, this balance reflects the familiar goal of reducing validation error through model parsimony.

Our models allow new video footage to be classified on a frame-by-frame basis, with some expected degree of accuracy. Linking this back into research objectives is the final step in the process. The seal classifier is intended to be used as a detection system. Even with a frame-specific false negative rate of 10%, no visits were missed entirely. An alarm system, triggered by *N* predicted presences in a sequence of *M* frames, is easily established, with *N* and *M* determined by balancing costs of false positives and negatives. Graphical displays such as Figure 2 convey this information in an easily digested way. Higher error rates prevent the use of the penguin behaviour classifier for the purpose it was intended for – replicating a human observer and calculating energy budgets – because certain classes of behaviour are poorly identified. However, surface behaviour was nearly perfectly distinguished from diving behaviour, and deep and shallow/subsurface dives were also well differentiated. These distinctions hold practical value, and also limit the amount of manual labelling that must be done.

Deep learning holds enormous promise for automating the labelling of video data, a process that looks increasingly unsustainable with manual methods. Case studies such as the ones reported here play an important role in reporting successes and failures, and developing and disseminating best practices. Classification of ecological data is difficult. Limited time and other resources, remote locations, and rare or difficult-to-detect target species, serve to decrease sample sizes at the same time that variable background environments increase the necessary sample sizes for good classification. In these contexts full automation is perhaps, for the time being, unrealistic. Facilitating the process of manually annotating video datasets is both valuable and achievable. Video data has the great advantage that large datasets, in terms of numbers of images, are often collected relatively quickly. At 60fps, a one minute encounter with an animal provides 3600 images. This offers exciting opportunities for developing and testing deep learning approaches. Our study suggest that many applications may benefit from incorporating temporal information in video, where the goal remains to predict the class to which a particular frame or image belongs. We expect these models to be widely used and developed in the near future.

## Acknowledgements

The penguin data collection was financially supported by Homebrew Films and the Percy Fitzpatrick Institute of African Ornithology. All fieldwork was done under permission from the South African Department of Environmental Affairs (permit nos RES 2015/38 and RES 2016/100) and Cape Nature (permit no. AAA007-00209-0056). We thank Cape Nature and the South African Department of Environmental Affairs for providing permission to carry out the study. The seal data collection was funded by the Scottish Government through the Marine Mammal Scientific Support Research Programme. ID is supported in part by funding from the National Research Foundation of South Africa (Grant ID 90782, 105782).

## Authors’ contributions

All authors conceived the work together. RH collected and annotated seal data, and provided feedback on model usability results. AM did the same for the penguin data. AC and ID developed the modelling approach. AC implemented the models and performed analyses. AC and ID wrote the paper. All authors contributed critically to the drafts and gave final approval for publication.

## Data accessibility

Code and analysis scripts are available online at https://github.com/alxcnwy/Deep-Neural-Networks-for-Video-Classification. A subset of seal and penguin video recordings, manual annotations, and results have been stored on Zenodo: https://doi.org/10.5281/zenodo.3842040.

## Notes

https://doi.org/10.5281/zenodo.3842040

https://github.com/alxcnwy/Deep-Neural-Networks-for-Video-Classification

### Competing Interest Statement

The authors have declared no competing interest.

