## Supplementary material 1 for "Frame-by-frame annotation of video recordings using deep neural networks"

#### A Examples and summaries of datasets

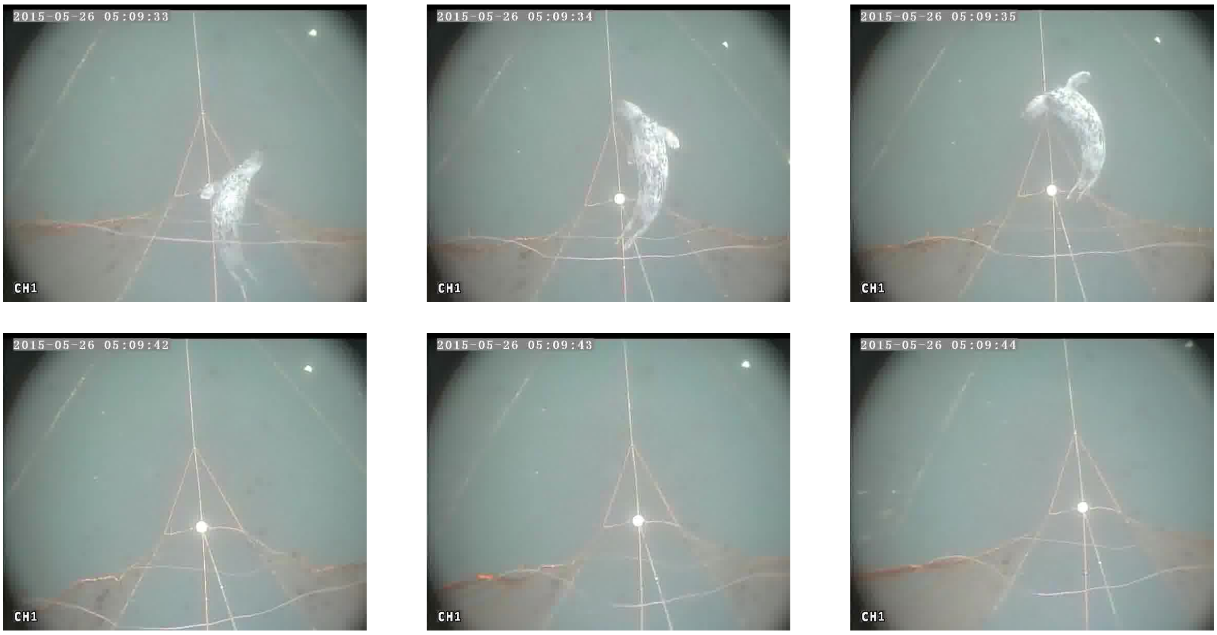


Figure A.1: Frame sequences showing a seal entering a net (top row) and typical background conditions in the absence of a seal (bottom row).

| | Descent | Bottom | Ascent | Shallow | Subsurface | Search |
|---|---|---|---|---|---|---|
| Train | 6030 | 4297 | 6394 | 7557 | 3663 | 14017 |
| Validation | 654 | 269 | 769 | 2317 | 666 | 2493 |
| Test | 524 | 244 | 733 | 278 | 168 | 1649 |

Table A.1: Penguins dataset class distribution across train, validation and test splits

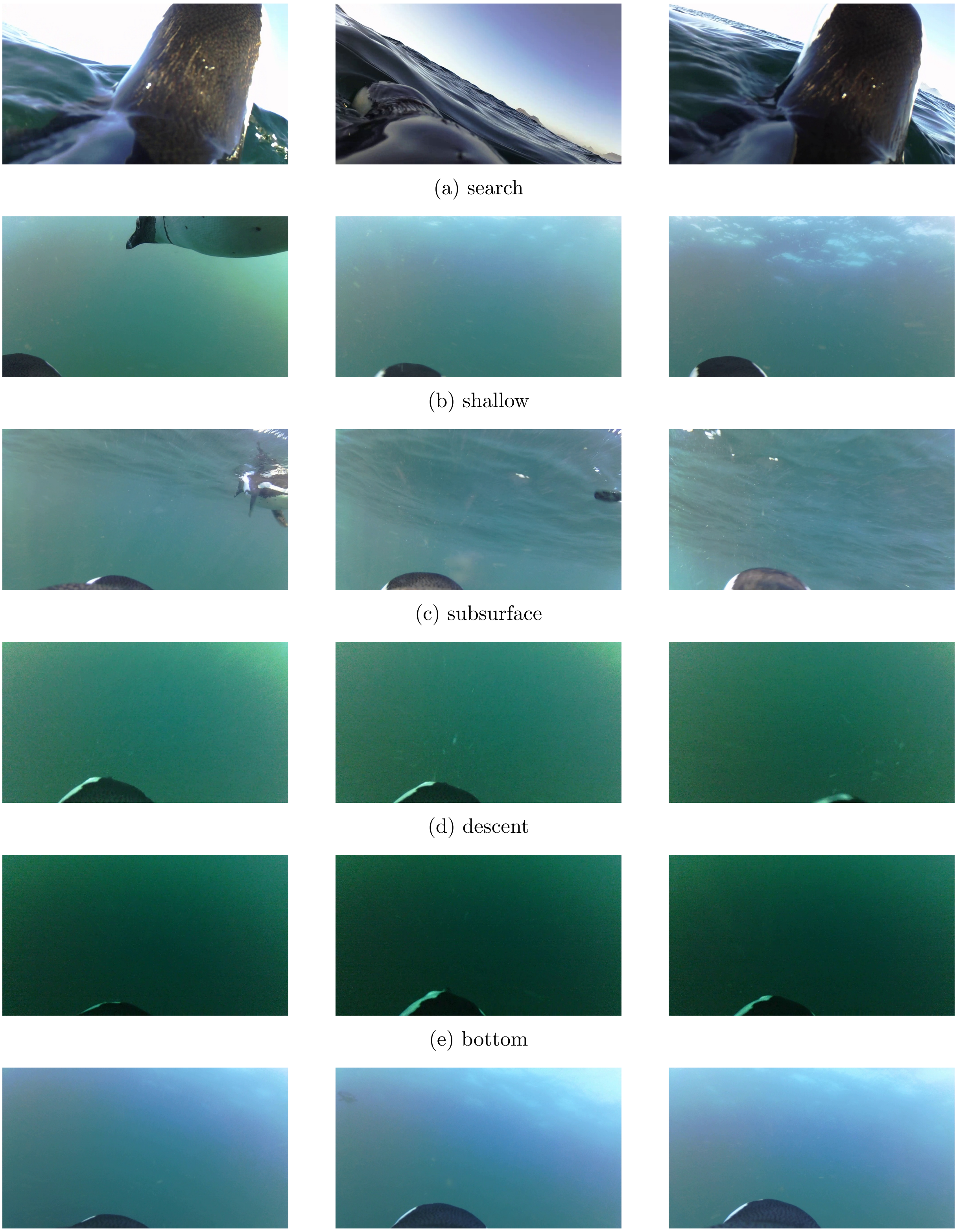

(a) search

(b) shallow

(c) subsurface

(d) descent

(e) bottom

(f) ascent

Figure A.2: Frame sequences showing each penguin behaviour class.

### B Further results

| | Precision (%) | | Recall (%) | | Predicted (Video) | | | | | |
|---|---|---|---|---|---|---|---|---|---|---|
| Observed | Video | Image | Video | Image | Dsc | Bot | Asc | Sha | Sub | Sea |
| Descent | 86 | 69 | 70 | 66 | 452 | 12 | 0 | 57 | 0 | 3 |
| Bottom | 57 | 43 | 62 | 53 | 88 | 139 | 17 | 0 | 0 | 0 |
| Ascent | 79 | 74 | 97 | 91 | 7 | 72 | 566 | 67 | 0 | 2 |
| Shallow | 63 | 62 | 46 | 34 | 93 | 0 | 0 | 176 | 3 | 6 |
| Subsurface | 48 | 48 | 96 | 83 | 2 | 0 | 0 | 80 | 81 | 5 |
| Search | 100 | 99 | 99 | 99 | 2 | 0 | 2 | 3 | 0 | 1641 |

Table B.1: Test set classifier accuracy within each behaviour for best video- and image-based penguin behaviour classifier, and a confusion matrix crosstabulating actual and predicted class membership in the best video model. Diving and surface activity is nearly perfectly discriminated, and including temporal information improves either precision or recall for most dive phases.

| | | | | |
|---|---|---|---|---|
| Accuracy (Test) | 89.4% | 89.2% | 89.1% | 89.1% |
| Precision (Test) | 100% | 99.4% | 100% | 97.6% |
| Recall (Test) | 83.9% | 83.9% | 83.3% | 84.9% |
| Accuracy (Validation) | 96.3% | 95.9% | 95.7% | 93.7% |
| Accuracy (Train) | 95.4% | 95.4% | 95.3% | 95.2% |
| Sequence Length | 5 | 5 | 3 | 1 |
| Architecture | RCNN | RCNN | RCNN | IMAGE |
| Train Duration (seconds) | 665 | 558 | 974 | 264 |
| Pretrained Model Name | resnet50 | resnet50 | resnet50 | resnet50 |
| Sequence Model | LSTM | LSTM | LSTM | NONE |
| Sequence Model Layers | 2 | 2 | 2 | NA |
| Layer 1 Size | 128 | 512 | 512 | 128 |
| Layer 2 Size | 128 | 512 | 512 | 512 |
| Layer 3 Size | 256 | 256 | 128 | 128 |
| Model Param. Count | 1 410 818 | 8 000 258 | 7 541 122 | 632 490 |

Table B.2: Classification accuracy and parameter settings for three best video models and best image model (detecting seals).

| | | | | |
|---|---|---|---|---|
| Accuracy (Test) | 85.4% | 84.0% | 84.2% | 80.5% |
| Precision (Test) | 85.4% | 84.0% | 84.2% | 80.5% |
| Recall (Test) | 87.6% | 87.6% | 85.5% | 82.8% |
| Accuracy (Validation) | 82.6% | 82.4% | 81.0% | 81.5% |
| Accuracy (Train) | 90.0% | 88.9% | 94.4% | 88.7% |
| Sequence Length | 20 | 20 | 20 | 1 |
| Architecture | RCNN | RCNN | RCNN | Image |
| Train Duration (seconds) | 570 | 663 | 1419 | 190 |
| Pretrained Model Name | vgg16 | vgg16 | vgg16 | resnet50 |
| Num. Features | 512 | 512 | 512 | 2048 |
| Sequence Model | SimpleRNN | SimpleRNN | LSTM | None |
| Sequence Model Layers | 2 | 2 | 2 | – |
| Layer 1 Size | 256 | 256 | 256 | 128 |
| Layer 2 Size | 128 | 512 | 512 | 256 |
| Layer 3 Size | 256 | 256 | 256 | 256 |
| Model Param. Count | 903 559 | 3 214 087 | 4 985 863 | 362 887 |

Table B.3: Classification accuracy and parameter settings for three best video models and best image model (classifying penguin behaviour).

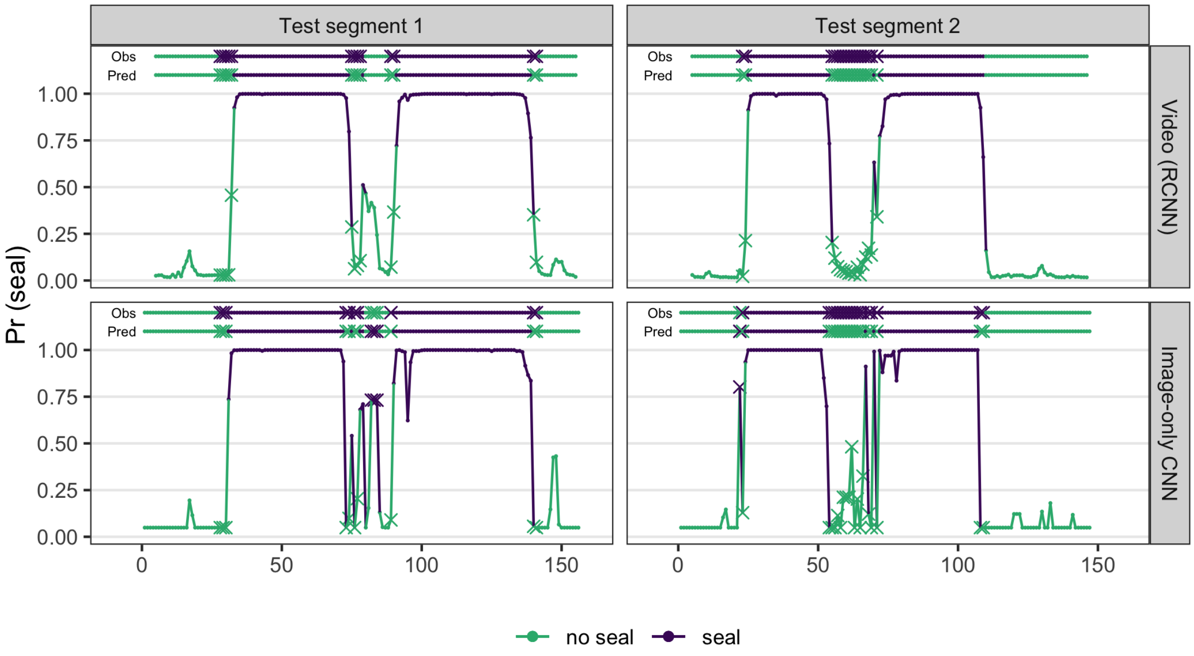


Figure B.1: Predicted probabilities of seal activity in salmon nets, with misclassifications plotted as crosses. Observed and predicted classes are plotted above the probabilities, using the same notation. Apart from one false negative (segment 2, frame 22–108), all incorrect classifications are at the beginning and end of visits, where only a small part of the seal may be in view. All visits are clearly identified.
